# Fast scanning small angle X-ray scattering of hydrated biological cells

**DOI:** 10.64898/2025.12.05.692524

**Authors:** Boram Yu, Mangalika Sinha, Rita Mendes Da Silva, Ulrike Rölleke, Manfred Burghammer, Sarah Köster

## Abstract

Due to their high penetration depth, X-rays enable us to obtain information from the interior of whole, unsliced cells. Scanning small angle X-ray scattering (SAXS), in particular, reveals real-space images in dark field representation as well as structural information in reciprocal space. However, obtaining information on anisotropy and orientation from cells in an aqueous, close-to-physiological environment remains challenging. Here, we extend the recently introduced fast scanning SAXS mode with short exposure times of few milliseconds to such hydrated samples by combining a newly developed, X-ray compatible microfluidic sample chamber and innovative data analysis that includes an effective noise-filtering method. This strategy enables the systematic analysis of radiation damage by quantifying the SAXS signal. Our results demonstrate that scanning SAXS can be used to obtain intracellular information of fixed-hydrated cells and the approach may in the future be applicable to living cells as well.

**Synopsis:** Fast scanning SAXS on biological cells in aqueous environment reveals intracellular anisotropy and orientation and allows for systematic assessment of radiation damage caused by the measurements.

## 1 Introduction

The mechanics, dynamics and function of biological cells are to a great extent determined by intracellular structures and imaging these nanometer- and micrometer-sized objects in a highly resolved manner has therefore become an essential approach for cell biology. The best-established methods for imaging of biological cells are electron and fluorescence microscopy, both of which have their distinct strengths. Electron microscopy (EM), in particular with the recent development of cryo EM enables extremely high spatial resolution of down to 1.55 Å, albeit only of static samples that are typically chemically fixed and dehydrated or even cryo-preserved (Fromm *et al*., 2023; Nogales & Mahamid, 2024). Fluorescence microscopy can be applied to samples in aqueous environment and even living cells. Super-resolution methods, such as MINFLUX now achieve nanometer resolution (Balzarotti *et al*., 2017; Schmidt *et al*., 2021).

A third, complementary probe for imaging are X-rays (Larabell & Nugent, 2010; Hémonnot & Köster, 2017; Weinhardt *et al*., 2019). Due to the small wavelength, and thus inherently high spatial resolution, and the high penetration depth, whole, unsliced cells can be imaged at high resolution. The electron density is imaged directly, thus avoiding the need to specific labels. Among a large variety of different X-ray imaging techniques, scanning small angle X-ray scattering (SAXS) with a focused beam provides real-space information at moderate resolution as well as reciprocal-space information at high resolution (Fratzl *et al*., 1997). Therefore, this method is employed to obtain multi-scale information of hierarchical structures of biological samples (Fratzl, 2002; Weinhausen *et al*., 2012). When first introduced, scanning SAXS was employed to investigate comparatively radiation-stable biological materials, such as wood (Fratzl *et al*., 1997), bone (Rinnerthaler *et al*., 1999), and teeth (Tesch *et al*., 2001). More recently others and we also applied it to radiation-sensitive biological cells (Weinhausen *et al*., 2012; Müller *et al*., 2010; Bernhardt *et al*., 2016; Bernhardt *et al*., 2017; Reichardt *et al*., 2020). Such studies are challenging, as the electron density contrast between the cells and the surrounding aqueous solution is weak and the signal easily gets lost due to radiation damage.

In the past, cells were studied in freeze-dried (Weinhausen *et al*., 2012; Wilke *et al*., 2012; Bernhardt *et al*., 2016; Bernhardt *et al*., 2017), frozen-hydrated (Priebe *et al*., 2014), fixed-hydrated, and even living conditions (Weinhausen *et al*., 2014; Priebe *et al*., 2014; Bernhardt *et al*., 2017). Typically, the total scan time for these studies extends from approximately 15 minutes to over one hour per single cell, depending on scan step sizes and exposure time. More recently, thanks to the high photon flux and consequently shorter exposure times after the EBS upgrade at ESRF, fast, continuous scanning enabled us to collect data from several hundreds of freeze-dried cells during a total experiment time of about 10 hours (Cassini *et al*., 2020). Dark field contrast images could be analyzed, including semi-automated cell segmentation of a large number of cells and *q*-dependent intensity profiles.

So far, fast scanning SAXS was constrained to freeze-dried samples, because the low signal-to-noise ratio (SNR) associated with the short exposure times is even lower in hydrated samples. In earlier studies employing conventional, slower scanning, data obtained from fixed-hydrated or living cell samples, enabled the analysis of dark field contrast images and *I*(*q*) profiles (Weinhausen *et al*., 2014) and spatial structural information, such as the extent of anisotropy (Bernhardt *et al*., 2017). However, derivation of meaningful structural information from fast scanning SAXS measurements on hydrated samples remains limited by low SNR.

Here, we present a strategy for imaging intracellular structures in fixed-hydrated samples by fast scanning SAXS including (i) a novel design for an X-ray compatible microfluidic chamber featuring continuous buffer flow and a minimized water layer thickness and (ii) effective noise-filtering method to extract meaningful information. We demonstrate that structural information, such as local orientation and anisotropy, can be obtained and we systematically assess the influence of radiation damage on the SAXS signal. Thus, our approach provides suitable measurement parameters for fast scanning SAXS on biological cells in aqueous environments.

## 2 Materials and methods

### 2.1 Sample preparation

We use SW-13 cells, derived from adrenocortical carcinoma (Leibovitz, 1973), stably transfected with fluorescence keratin hybrid expression (HK8-CFP, HK18-YFP) (Strnad *et al*., 2002; Windoffer *et al*., 2004). These cells are cultured in high glucose (4.5 gL^−1^) Dulbecco’s Modified Eagle’s Medium (DMEM, D6429; Merck KGaA, Darmstadt, Germany) supplemented with 10 %(v/v) fetal bovine serum (FBS, 10270-106, Gibco), and 1 % (v/v) 100 UmL^−1^ penicillin and 0.1 gL^−1^ strep-tomycin (pen-strep, 15140122; Gibco). Following the thawing process, the cells are incubated at 37°C in a water-saturated atmosphere with 5 % CO_2_.

Silicone nitride membranes (SiN, frame size 10 mm *×* 10 mm, frame thickness 200 *µ*m, window size 1.5 mm *×* 1.5 mm, window thickness 1 *µ*m; Silson Ltd, Warwickshire, UK) are plasma-activated at 50 W for 40 s (Zepto, Diener electronic GmbH & Co. KG, Ebhausen, Germany) to render the surface hydrophilic. Subsequent to the plasma-activation, the membranes are immersed in ultrapure water and undergo UV treatment for 30 minutes for sterilization. The sterilized membranes are then positioned on a 12-well cell culture plate, with the flat side oriented upwards. A solution of 40 *µ*L of 5 % (v/v) fibronectin (F1141, Merck KGaA) in phosphate buffered saline (PBS, D8537, Merck KGaA) is applied to the membrane to enhance cell adhesion and incubated at 37°C for one hour.

Once approximately 80 % confluency of the cultured cell layer is reached, the cells are detached by adding 0.25 % (v/v) trypsin with 0.02 % (w/v) EDTA (P10-020500, PAN-Biotech GmbH, Aidenbach, Germany) in PBS, and thereafter suspended in fresh culturing medium. Following this step, the cells are seeded onto the membrane at an initial concentration of 1.5 *×* 10^5^ cells/mL, or 6.25 *×* 10^5^ cells/cm^2^, i.e., 1.5 mL in each well in a 12-well cell culture plate. After a 24-hour incubation, the membranes are washed with PBS and fixed for 15 minutes with a 4 % methanol-free formaldehyde solution (28906, Thermo Fisher Scientific Inc., Darmstadt, Germany) in PBS, followed by three rinses with PBS (Thavarajah *et al*., 2012).

Visible light phase contrast and fluorescence images of the membranes with fixed cells are recorded using an inverted microscope (IX83 Olympus, Hamburg, Germany), equipped with a UCPlanFLN 20*×* objective (0.7 NA, Olympus) and UPLFLN 40*×* objective (0.75 NA, Olympus). For fluorescence imaging, an IX3-FGFPXL filter set (excitation filter: 460-480 nm, emission filter: 495-540 nm, dichroic mirror: 490 nm; Olympus) is employed. Thereafter, the fluorescence images are sharpened by Fourier filtering (Petrou & Petrou, 2010) using the python tool box, scikit-image (Van der Walt *et al*., 2014). The membranes are individually stored in PBS with 1 % (v/v) pen-strep and transported to the ESRF.

### 2.2 The microfluidic chamber

Master wafers for the polydimethylsiloxane (PDMS, Sylgard 184, Dow Corning, Midland, MI, USA) structures including the flow channels are fabricated by standard photolithography. Briefly, SU-8 negative photoresist (SU-8 3050, MicroChem, Newton, MA, USA) is spin coated onto a silicon wafer (2-inch, MicroChemicals, Ulm, Germany) to a height of 210 *µ*m. To reach this height, the process has to be repeated twice. The resist is exposed to UV light through a mask containing the structure (see Fig. S1a in the Supplementary Information) and developed. The wafers are coated by fluorosilane (1H,1H,2H,2H-perfluorooctyltriethoxysilane, 97 %, AB 104055; abcr GmbH, Karlsruhe, Germany) and PDMS replicas are produced from the wafers. One PDMS replica needs to be thick enough (10 mm) to distribute the pressure generated by clamping the edges with the metal frame to the SiN membranes, thus providing a good seal. The second replica is produced as thin as possible (5 mm) so as to position the sample as close to the X-ray aperture as possible. After curing, the PDMS is removed from the wafer, holes are punched for the inlet, outlet (diameter 0.75 mm) and X-ray observation window (diameter 3.5 mm) (see Fig. S1b in the Supplementary Information). A schematic of the chamber preparation is shown in Fig. S2a-d in the Supplementary Information.

Just before the experiment, the PDMS replicas are plasma-activated at 15 W for 60 - 80 s (Tergeo, PIE Scientific LLC, Union City, CA, USA). The SiN membrane with the cells is rinsed with ultrapure water to remove salt and to avoid crystal formation during scanning. The water on the side opposite of the cells is completely blotted off with filter paper. The SiN membranes are placed above the observation windows in the plasma-activated PDMS replicas, and the two halves are assembled under a stereomicroscope (Wild M3Z, Leica Microsystems GmbH, Wetzlar Germany). Tubings (inner diameter: 0.38 mm, outer diameter: 1.09 mm, polyethylene; Becton, Dickinson and Company, NJ, USA) filled with degassed PBS are inserted into the chamber, and all components are clamped by the metal frames. The assembly process, including the sample preparation process explained in Sec. 2.1, is illustrated in Fig. S2e-j in the Supplementary Information.

When fully assembled, the sample chamber is composed of two SiN membranes, one of which the cells are grown, and the other one with an SU-8 spacer of a thickness of 20 *µ*m (Silson Ltd) on its flat side (see Fig. 1a and Fig. S1c in the Supplementary Information for more details), thereby enabling liquid flow over the window region. The two SiN membranes are arranged in such a manner that their flat sides face each other. The sandwiched membranes are encased by two pieces of PDMS with a flow channel structure of 1.5 mm width and 30 mm length, including a 10 mm square positioned centrally, wherein the membranes are placed (see Fig. S1a in the Supplementary Information). The components are clamped by two metal frames (see Fig. S3a in the Supplementary Information), which ensure leak-tightness during prolonged operation. The assembled chamber is inserted in a dedicated chamber holder compatible with the X-ray sample stage described in Sec. 2.3 (details are provided in Fig. S3 b-c in the Supplementary Information). The resulting chamber has an operation time over 24 hours and can in principle be used with flow rates of more than 1000 *µ*Lh^−1^. A syringe pump (base and single module, Cetoni GmbH Korbussen, Germany) is used to control the volume flow and we adjust the flow rate between 20 *µ*Lh^−1^, which corresponds to 0.18 mms^−1^, to 1000 *µ*Lh^−1^, which corresponds to 9.26 mms^−1^ for the given channel geometry.

**Figure 1.**
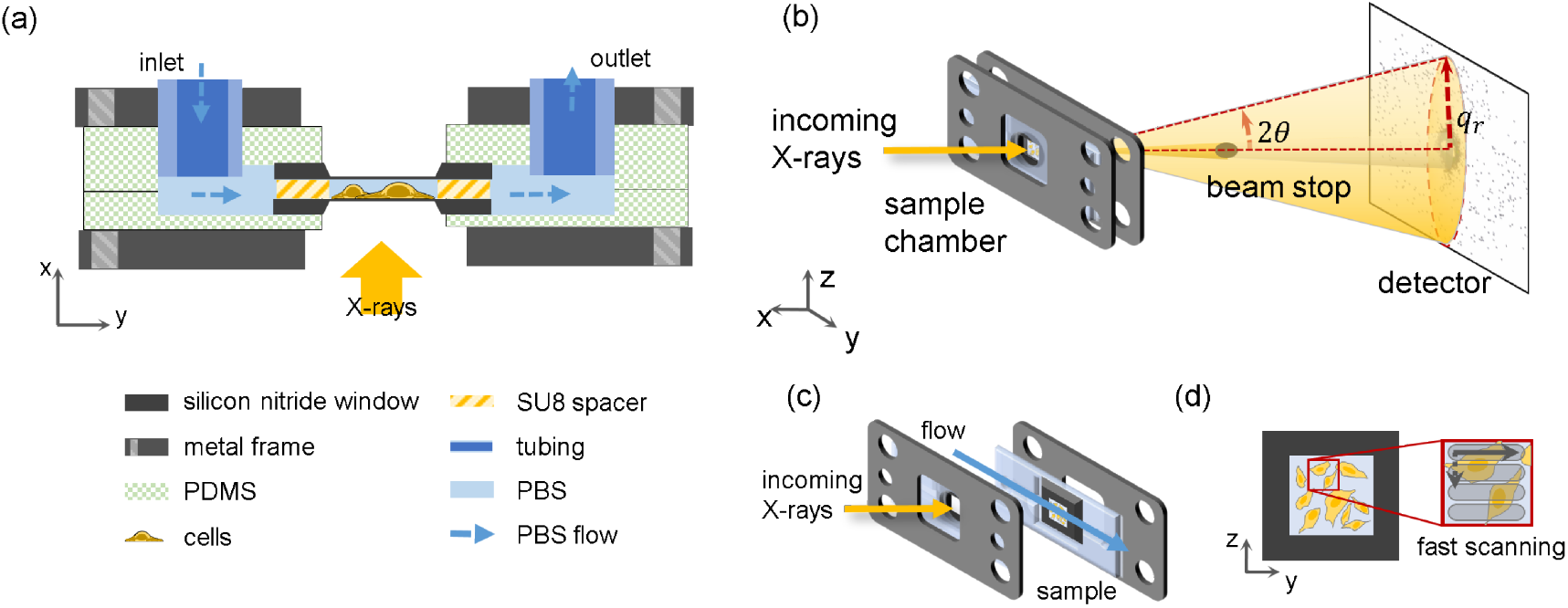
Experimental setup. (a) Schematic side view of the dedicated flow chamber (see legend for the individual components). Two SiN membranes are separated by 20 *µ*m using SU-8 spacers. This “sandwich” is enclosed by PDMS layers and clamped by metal frames. The buffer flow is represented by blue dashed arrows. (b) Schematic of the SAXS experiment. The sample (fixed-hydrated cells on a SiN membrane) is scanned through the focused X-rays. The beamstop blocks the direct beam, and the two-dimensional detector records the scattered X-rays. The red dashed lines represent the scattering angle 2*θ* between the incoming beam direction and the scattered X-rays. (c) The flow within the microfluidic chamber is oriented horizontally in perpendicular direction to the incoming beam, thus along the *y*-axis. (d) Representation of the scanning scheme; the SiN membrane is moved along the *y*- and *z*-axes to obtain a raster scan of the selected region with the specified step numbers and sizes.

### 2.3 Scanning SAXS

The scanning SAXS experiments are performed at the nano-branch of beamline ID13 of the European Synchrotron (ESRF) in Grenoble, France. A sketch of the experiment geometry is shown in Fig. 1b. The X-ray beam from the undulator is monochromatized by a Si-111 channel-cut monochromator to an energy of 15 keV. The beam is focused through multiple steps: The beam is first pre-focused by beryllium parabolic refractive lenses (Be-CRLs). It is then focused by multilayer Laue lenses (MLL) to a final size of 250 nm *×* 250 nm at a flux of 1.09 *×* 10^12^ photons/s.

At the end station, the focused beam undergoes conditioning by a square shaped 40 *×* 40 *µ*m^2^ PtIr aperture (order-sorting aperture, OSA) and is subsequently targeted at the sample. At the downstream side of the sample, a helium-filled flight tube is placed to reduce the air scattering and the beamstop is located right outside the exit window of the tube to block the direct beam. Scattered X-rays are recorded with an Eiger X 4M two-dimensional X-ray detector (2070 *×* 2167 pixels^2^, i.e., about 4 megapixels with a pixel size of 75 *µ*m *×* 75 *µ*m; Dectris, Baden, Switzerland). The distance from the sample to the detector is 1.89 m, as determined by measuring AgBeh powder in the empty chamber.

The assembled sample chamber is mounted on a piezo translation stage, which is installed on a hexapod. The hexapod is used for coarse positioning, while the piezo stage facilitates finer motion with a minimum step size of 20 nm; thus, this configuration enables fast scanning. The scanning is conducted along the *y*- and *z*-axes as shown in Fig. 1, which form a vertical plane to the beam path. Photographs of the end station after mounting the chamber are shown in Fig. S4 in the Supplementary Information. The mounted sample is aligned with the beam using an on-axis microscope which can be moved in and out of the beam.

During the measurement, the step sizes for the *y*- and *z*-axes are selected with respect for the beamsize: similar to the beam size (200 nm *×* 200 nm), which yields the highest spatial resolution but also higher radiation damage, four times the beam area (500 nm *×* 500 nm) to minimize radiation damage at the expense of resolution, or asymmetrically (200 nm *×* 500 nm), which balances radiation damage and spatial resolution. The exposure time is set just above the detector’s lower limit (1.34 ms) and thus long enough to collect a sufficient signal while minimizing radiation damage. The radiation dose *D* can be estimated by

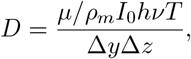

where *µ/ρ*_*m*_ = 1.60 cm^2^g^−1^ is the ratio between mass attenuation coefficient and mass density of the cellular material (Berger *et al*., 2010), *I*_0_ is the photon flux of the incoming beam, *hν* is the photon energy, *T* is the exposure time per scan point, and Δ*y* and Δ*z* are the step sizes (Howells *et al*., 2009; Weinhausen *et al*., 2012). The measurement parameters and estimated radiation doses are listed in Table 1.

**Table 1:** Measurement parameters and calculated radiation doses.

| $\Delta y/\mu\text{m}$ | $\Delta z/\mu\text{m}$ | exposure time/ms | dose/$10^7\text{Gy}$ |
| --- | --- | --- | --- |
| 0.2 | 0.2 | 2 | 2.09 |
| 0.2 | 0.5 | 2 | 0.84 |
| 0.2 | 0.5 | 5 | 2.09 |
| 0.5 | 0.5 | 2 | 0.33 |
| 0.5 | 0.5 | 5 | 0.84 |

In order to analyze the decay of the ordered structures in response to the radiation dose, scanning SAXS is conducted in a single area multiple times with the same scanning parameters.

### 2.4 Data analysis

Each scan consists of 40,000 to 250,000 individual scattering patterns. The scattering intensities, along with their corresponding positions, are used to calculate a dark field contrast image, where the total scattering intensity is plotted on a color scale (see Fig. 3a) (Bunk *et al*., 2009; Weinhausen *et al*., 2012). The background and foreground regions are segmented based on the dark field images. To account for beam variations during the extended scans (15 - 20 minutes per scan), the background region is determined by automated cell segmentation via adaptive thresholding (ACSAT, (Shen *et al*., 2018)). ACSAT is a cell segmentation method that employs global and local thresholding iteratively. In each iteration, the algorithm detects the foreground regions at various threshold values and creates regions of interest (ROIs) containing a cell via a morphological operation. It then selects the minimum threshold value that leads to the maximum number of ROIs. This process is conducted for the entire image (global thresholding) first, and is then repeated in a rectangular ROI (local thresholding). During this process, any detected ROIs smaller than *A*_*min*_ are filtered and removed. For choose *A*_*min*_ = 500 pixels for scans with 500 *×* 500 nm^2^ step size; *A*_*min*_ = 750 pixels for 200 *×* 500 nm^2^; *A*_*min*_ = 1000 pixels for 200 *×* 200 nm^2^. The cell mask created by ACSAT is shown in Fig. S5b in the Supplementary Information. Remaining noise is filtered out using a histogram-based Gaussian filter, which removes pixels whose intensity exceeds 3 standard deviations (3*σ*) from the Gaussian distribution of the masked regions’ intensities. These filtered pixels and the cell regions are dilated by 3 pixels with a circular structure element, and the background region is defined as non-selected pixels. The background signal, which is used to correct the data in a row-by-row manner, is defined by the averaging the scattering patterns of all pixels of the background region in a single row.

The local orientation, indicating the average orientationof all objects at the measurement position, is obtained by calculating the circular mean and the circular variance (Allen & Johnson, 1991; MacArthur & Thornton, 1993) of the azimuthal intensity profile, which correspond to the orientation and anisotropy, respectively. By plotting the orientation and anisotropy values at their corresponding positions, an orientation map as well as an anisotropy map are obtained (see examples in Fig. 3b, c). Following these analyses, the ordered intracellular structures, in our case, e.g., bundles of keratin filaments, are visible, and the decay of the order with increasing radiation dose is analyzed. A detailed graphical description of the data analysis workflow is provided in Fig. S6 in the Supplementary Information.

The decay of the ordered intracellular structures as a response to the radiation is computed for individual cells. For this analysis, each individual cell in the background-subtracted dark field contrast image is annotated by marker-controlled watershed segmentation (Meyer, 1994). In order to annotate the cells individually, a distance map is calculated for a watershed segmentation. For separating adjacent cells, we make use of the fact that each cell contains one nucleus. A nuclei mask is created via morphological reconstruction. Morphological reconstruction is a feature extraction method based on pixel connectivity that preserves the original shape of the nuclei (Vincent, 1993). Applying the mask as the local minima for the watershed segmentation, all cells, including adjacent cells, are annotated individually. The final cell masks, the nuclei masks, and the cell annotation result are shown in Fig. S5b-d in the Supplementary Information.

Following the annotation step, anisotropy distributions of ordered intracellular structures of individual cells are calculated. As ordered structures are more anisotropic, pixels in a cytoplasmic region with high anisotropy values are selected. The cytoplasmic region is defined as the area that is included in the cell mask but excluded from the nuclei mask. To make this selection, an anisotropy distribution of the masked background region is plotted, and its 99-th percentile value is used as the thresholding value. To track changes in the anisotropy distributions induced by radiation, the distributions are depicted as violin plots against scan number. The difference in mean value between the first and the second distribution is set as the anisotropy decay caused by radiation damage and then compared against different radiation doses.

All data analysis is performed using MATLAB R2020a (The MathWorks, Inc., Natick, MA, USA) scripts, including the Image Processing Toolbox and functions of the Nanodiffraction toolbox presented in Ref. (Nicolas *et al*., 2017).

## 3 Results and discussion

We perform fast scanning SAXS in biological cells grown on SiN membranes and incorporated in a dedicated microfluidic flow chamber. Due to the short exposure time of few ms and the water layer surrounding the cells, as well as the low electron density contrast between water and cells, the collected SAXS data show a low SNR ratio, making the data analysis challenging (Bernhardt *et al*., 2017; Nicolas *et al*., 2019*b*; Reichardt *et al*., 2020). Here, we propose a novel approach to filter noise in the data that involves constraining the *q*-range based on a systematic analysis of dark field contrast images calculated from different *q*-ranges. This method allows for the extraction of structural information of intracellular components and we demonstrate the utility by quantitatively analyzing the impact of radiation damage to intracellular structures.

### 3.1 Optimization of the *q*-range for data analysis

In scanning SAXS, dark field contrast images are calculated by summing up the intensity values of all pixels across the entire detector area (Fratzl *et al*., 1997; Priebe *et al*., 2014) or just within a circular region around the primary beam (Bunk *et al*., 2009; Weinhausen *et al*., 2012). In many cases, the appropriate detector range is chosen by a qualitative inspection of the resulting dark field images. In our case, however, due to the fast scanning, short exposure times and, therefore, strong noise, such an approach is difficult. Instead, we choose an objective way to optimize the *q*-range for the dark field images, which is then used for further analysis of the SAXS data as well.

We calculate the magnitudes of the scattering vectors *q* as

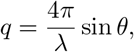

where *λ* is the wavelength of incoming X-rays, and *θ* is half the scattering angle (see Fig. 1b). For our setup the maximum *q*-value is 5.465 nm^−1^.

We determine dark field contrast images using various *q*-values ranging from 0.003 nm^−1^ to 5.465 nm^−1^ with 0.003 nm^−1^ intervals, resulting in 1,795 data points in total. The intensity values in the images are normalized to a range of [0, 1]. We compare the noise level of the individual images, defined by the mean deviation (MD, also called the mean absolute deviation) of a rectangular, featureless region of size *a × b*:

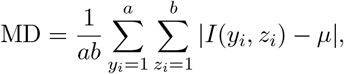

where the mean intensity is given by

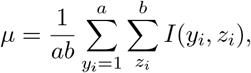

and *I*(*y*_*i*_, *z*_*i*_) is the intensity of the pixel (*y*_*i*_, *z*_*i*_) (Sari *et al*., 2012; Rajabi & Zirak, 2016). The mean deviation is more suitable than the standard deviation because it is less sensitive to extreme values. The MD values for the selected background region are plotted against *q* in Fig. 2a. As the MD is obtained from the normalized dark field images, it is indicative of the SNR of the image. The maximum of the MD curve *q*_1_ is chosen as the lower bound of the optimized *q*-range, because it corresponds to the minimum radius at which scattering signal from the sample is detected, and the minimum of the MD curve *q*_3_ as the upper bound, because here the scattering signal from the cells is sufficiently strong while the background noise is minimal. Before determining the minimum of the MD curve, it is smoothed with a window size of 38 data points, determined by the elbow method (Thorndike, 1953). The choice of the upper bound is illustrated by an image series for selected *q* values shown in Fig. 2c. Here, we show five images for comparison, which are selected as described in the caption. The cell contours are hardly visible when *q*_1_ is very small. For *q*_2_ = 0.177 nm^−1^, the cell contours are clearly visible, however, the *q*-range is still very small (see Fig. 2b). This indicates that most signal stemming from the cells is concentrated close to the center of the scattering pattern at small *q*-values. When *q*_3_ = 0.299 nm^−1^, corresponding to the minimum of the curve, the background area surrounding the cells becomes brighter than in the second image, indicating improved contrast. However, upon expanding the calculation region to *q*_4_ = 2.513 nm^−1^ and *q*_5_ = 5.465 nm^−1^, stripes along the fast scanning direction (*y*-axis in Fig. 1d) appear. These stripes are considered additive noise (see Fig. S7 in the Supplementary Information) because they are independent of the original signal. This suggests that the observed noise is caused by exterior sources, such as mechanical vibrations of the stage during scanning. These example dark field contrast images show that the noise level depends on the *q*-range considered for the calculation. For the work presented here, we determine *q*_opt_ to be *q*_3_ from the specific data set shown in Fig. 2 and use it for the analysis of all data sets.

**Figure 2.**
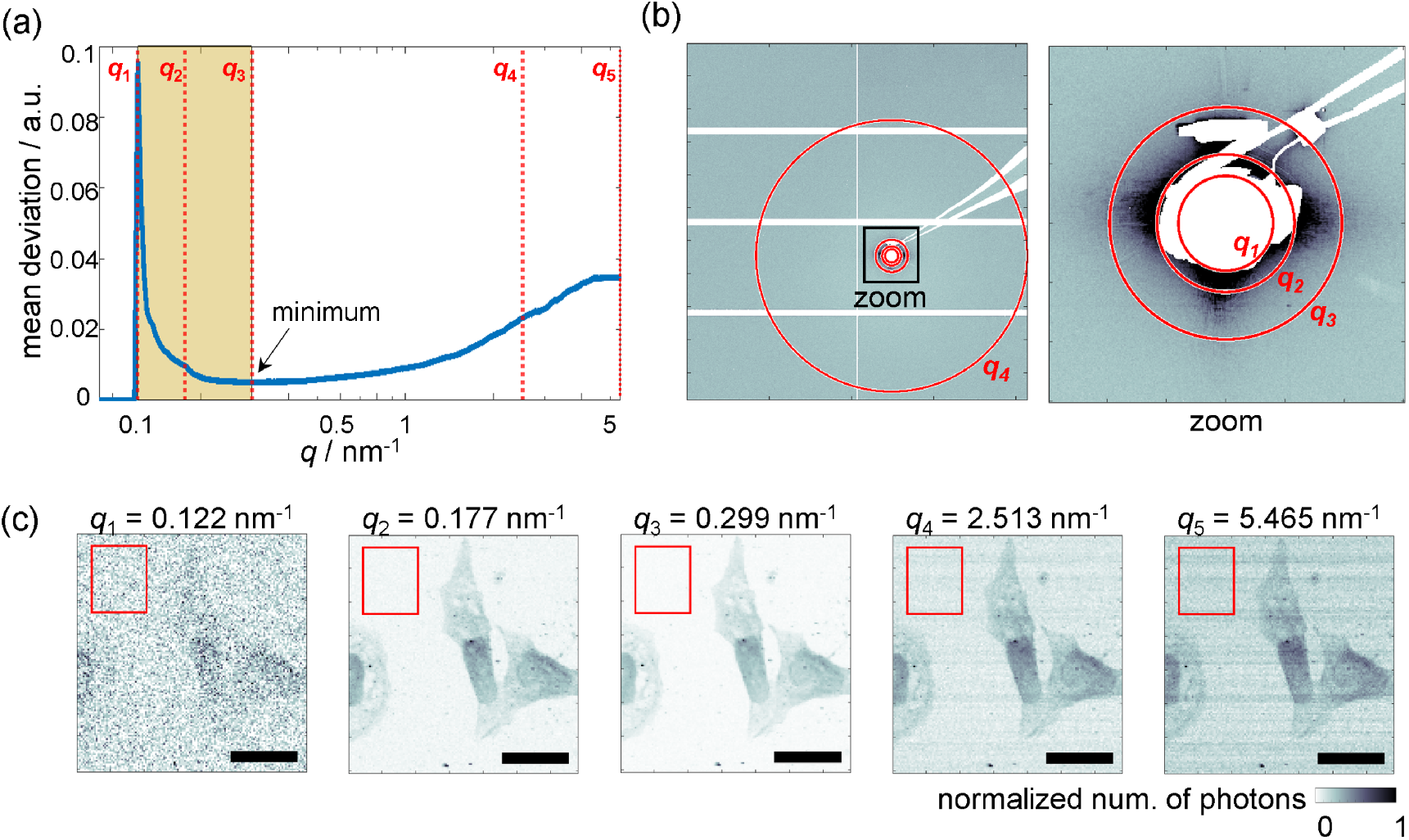
Optimization of the *q*-range. (a) The mean deviation of the background region (red boxes in panel c) is plotted against the magnitude of the scattering vector *q*. The vertical red dashed lines correspond to the selected *q* values in b and c. The chosen *q*-range (yellow), reaches from *q*_1_ to *q*_3_. (b) Summed scattering pattern from 40,000 individual patterns of the whole detector view (left) and an enlarged view (right). The beam stop and aperture edge artifacts are masked and shown in white. (c) Series of X-ray dark field contrast images obtained with varying maximum *q*-values *q*_*i*_ as shown in panel b; *q*_1_ is the minimum radius at which scattering signals appear, *q*_2_ is a circumscribed circle of the beam stop (ignoring neighboring artifacts), *q*_3_ is the minimum of the mean deviation curve in panel a, *q*_4_ corresponds to the maximum circle within the detector area and *q*_5_ corresponds to the whole detector area. In each case the minimum *q* value used for the calculation of the dark field image is *q*_1_. The scanning is performed with 0.5 *×* 0.5 *µ*m^2^ step size, and 5 ms of exposure. Scale bars are 20 *µ*m.

### 3.2 Analysis of the local orientation of cellular structures

Using the *q*-range from *q*_1_ to *q*_opt_, we further analyze the example data set shown on Fig. 2. A dark field contrast image of a slightly larger field of view is shown in Fig. 3a, on a logarithmic color scale. Distinct intensity differences are observed between the cytoplasmic and nuclear regions, with some variations in intensity within each region. As shown previously, these variations arise from inhomogeneous distributions of the cellular components, such as the cytoskeleton in the cytoplasm (Weinhausen *et al*., 2012), or different degrees of DNA compaction in the nuclei region (Hémonnot *et al*., 2016). Thus, the dark field contrast images provide real space information and inform us about the location of cellular structures such as the nucleus. However, in scanning SAXS, the scattering pattern recorded in each pixel contains additional reciprocal space information on the local structure within the cell. In particular, we analyze the local orientation and calculate orientation and anisotropy maps, as shown in Fig. 3b and c, respectively. Here, we restrict the analysis to the area within the white box shown in Fig. 3a.

**Figure 3.**
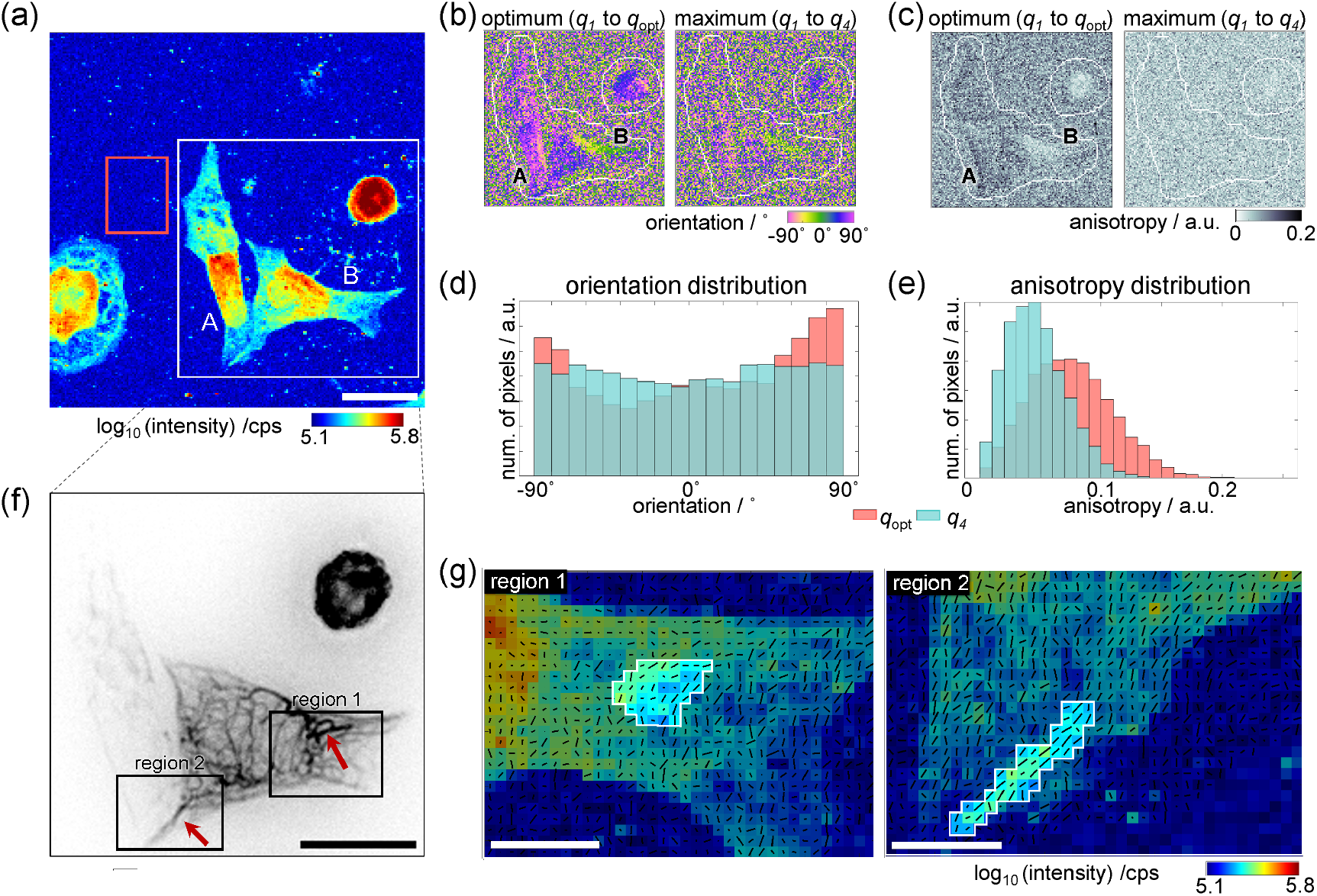
Orientation of cellular structures (a) Dark field contrast image of the same dataset as shown in Fig. 2c with a larger field of view. The red rectangular region in Fig. 2c is also added for comparison. (b) Orientation maps of the white rectangular region in panel a, calculated using different *q*-ranges: from *q*_1_ to *q*_opt_ (optimum range as discussed in Sec. 3.1) and from *q*_1_ to *q*4 (maximum range for local orientation analysis). (c) Corresponding anisotropy maps. The white outlines in panel b and c delineate the border of the background area used for the background subtraction. Histograms of (d) orientation and (e) anisotropy distributions of cells A and B as indicated in panels a-c. (f) Inverted gray scale epifluorescence image of the same cells, acquired prior to the microfluidic chamber assembly. The dark structures are distinct keratin bundles within the cell. (g) Dark field contrast images of the two regions 1 and 2 within the black boxes in panel f, where strongly ordered keratin structures are visible. The black lines in each pixel represent the orientation and anisotropy by their angle and length, respectively. Scale bars in panel a and f: 20 *µ*m, in panel g: 5 *µ*m.

We compare the local orientation for two different *q*-ranges, i.e., the optimum range from *q*_1_ to *q*_opt_, and the maximum range from *q*_1_ to *q*_4_, as shown in Fig. 3b, c. In panel b, an orientation of 0° corresponds to the horizontal direction. Note that for calculating the azimuthal intensity profile a perfect ring-shaped region needs to be considered (Bernhardt *et al*., 2016). We restrict our analysis of this example to the two cells marked by A and B in Fig. 3a-c as the corresponding fluorescence image (Fig. 3f) informs us that they are nicely spread out and attached to the substrate. The orientation and anisotropy maps in Fig. 3b, c clearly show that the cell outlines are more distinct when applying the optimal *q*-range, since the cellular material is both anisotropic and oriented, whereas the background is not. By contrast, if the *q*-range is increased, the signal is “diluted” by featureless background signal. The histograms in Fig. 3d, e display the orientation and anisotropy distributions of cells A and B. The orientation distribution of the optimum range (red) shows dominant distribution features at *±*90° (vertical direction in the image), while that of the maximum range shows a uniform distribution over all angles. In the case of the anisotropy, the distribution shifts towards the lower values when increasing the *q*-range, in agreement with our qualitative assessment regarding Fig. 3c. Taken together, these quantitative comparisons highlight the importance of choosing the optimal *q*-range for detecting the weak cellular SAXS signal and support our method for determining *q*_opt_ as discussed above. Notably, the anisotropy is more affected by the *q*-range than the orientation. The anisotropy is obtained by the circular variance of the azimuthal intensity profile, while the orientation is obtained from the circular mean. Increasing the *q*-range beyond the *q*_opt_ introduces noise to the azimuthal intensity profile at all angles, and the extent of this noise grows with *q*. As the *q*-range is increased, the circular mean remains stable until the noise level exceeds a certain threshold. However, the circular variance is directly affected by the added noise, and thus it changes promptly. To illustate this argument, a simulation comparing the sensitivity of the circular mean and the circular variance on the noise level is shown in Fig. S8 in the Supplementary Information.

A closer comparison of the local orientation results of the optimum *q*-range with the dark field contrast image reveals a correlation between orientation and cell shape. Cell A is elongated in the vertical direction (*±*90°, purple). The right side of cell B is elongated in a predominantly horizontal direction (0°, green). The anisotropy is increased in certain cellular regions and high anisotropy values are found at the cell contours, whereas low values are found in the nuclear regions in both cells, coinciding with high electron density found in the dark field image (Fig. 3a). We thus hypothesize the oriented, anisotropic signal to stem from cytoskeletal structures within the cytoplasm.

To investigate this hypothesis further, we compare a fluorescence micrograph (Fig. 3f, inverted gray scale, dark regions show labeled keratin bundles (Weinhausen *et al*., 2012)) of the example region with the SAXS data. We focus on two regions 1 and 2 (black boxes), where the strongly ordered keratin network is visible in the fluorescence image. Region 1 contains a thick keratin bundle forming a loop-like structure and this region is marked by the white outline in Fig. 3g, left. The black lines overlaid to the dark field image show the orientation and anisotropy by their angle and length, respectively and we can follow the loop structure here as well. Region 2 shows a cellular extension that contains a thick keratin bundle and the SAXS data (Fig. 3g, right) show increased electron density (dark field image) and aligned, anisotropic signal (black lines). More examples are shown in the Supplementary Information in Fig. S9 and S10.

### 3.3 Radiation damage from repeated imaging of cells

Cellular structures are generally very sensitive to the high radiation dose input by a focused X-ray beam. In order to investigate how this radiation damage influences the anisotropic, oriented signal stemming from cytoskeletal structures, we scan the same sample region multiple times with the same scanning parameters, i.e., step size and exposure time. An example series of dark field contrast images of the first, second, and fifth scan of one such a region is shown in Fig. 4a. Visual inspection for comparison of the first and the second dark field image shows changes in the cytoplasmic regions, where the intensity values are slightly decreased, yet the cell shapes are still intact and clearly distinguished. However, the cell shapes are deteriorated in the fifth scan image. In contrast, the nucleus appears more resistant to the radiation and the shapes of the nuclei are still clearly visible in the fifth scan.

**Figure 4.**
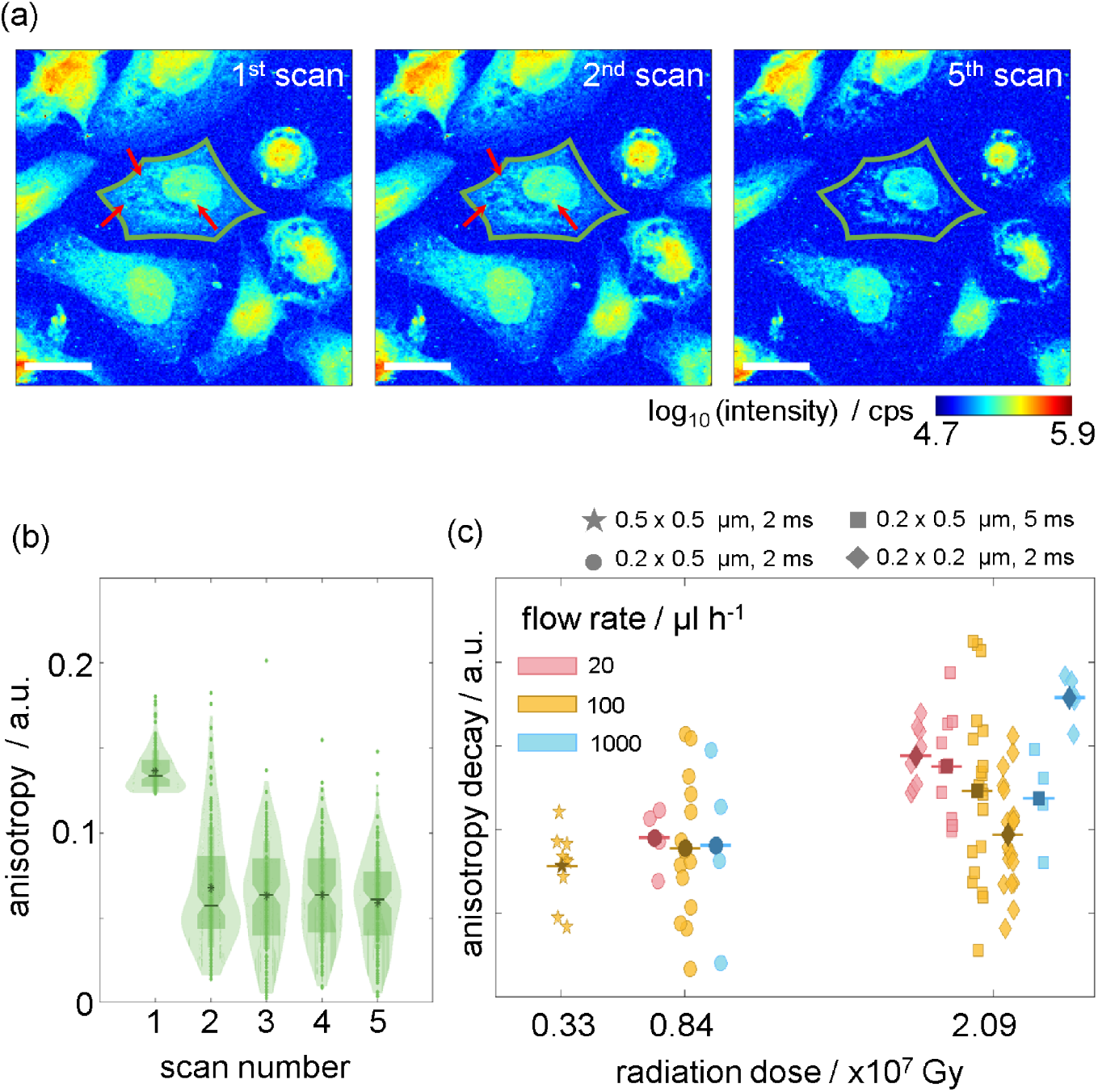
Analysis of radiation damage. (a) First (left), second (center) and fifth (right) dark field contrast image obtained from multiple scans of the same region. Each cell in the first scan is labeled individually, represented by different colors in Fig. S6 in the Supplementary Information. Scale bars: 20 *µ*m. (b) Anisotropy distribution of a single example cell that is labeled in green in panel a plotted against the scan number. In the cytoplasm region of individually labeled cells, while excluding the nuclear region, only pixels with anisotropy values greater than the 99th percentile of the anisotropy distribution of the background region are used. (c) Degree of anisotropy decay from the first to the second scan of individual cells using different measurement conditions (step size and exposure times, see legend above the figure). The color of each symbol corresponds to the specified buffer flow rate. Darker symbols show the mean values for each respective condition.

As the cellular material persist on the substrate after the first scan and even after four scans, we apply the analysis explained in section 3.2 to these repeated scans. In Fig. 4b, we show the anisotropy distributions of the single cell outlined in green in Fig. 4a plotted against the scan numbers. A strong decrease in anisotropy is visible only between the first and the second scan, thereafter we do not observe considerable changes anymore. This result indicates that most ordered structures are destroyed during the first scan already. This is remarkable, as the dark field image calculated from the second scan (Fig. 4a, center) looks similar to the one from the first scan (left), with the cytoplasmic regions retained.

Considering the strong decrease in anisotropy after the first scan, we quantify the extent of radiation damage of the ordered structures using the anisotropy decay defined as the difference in mean value between the first and the second distribution. The anisotropy decay values of individual cells measured using different scanning step sizes and exposure times are plotted against the corresponding radiation dose, as shown in Fig. 4c. Different scanning parameter sets are denoted by different symbols, and the flow rates are represented by different colors. One notable result is that as the radiation dose increases, the mean as well as the variance of anisotropy decay distributions increases. The large variance is due to the natural cell–to–cell variability. Given that anisotropy decay cannot be negative, the possible anisotropy decay value is equivalent to or smaller than the initial anisotropy values. It follows that some cells whose mean anisotropy values are initially small and completely damaged afterward result in small anisotropy decay values. Conversely, other cells initially showing large anisotropy values result in various values depending on the extent of the damage. A higher radiation dose yields a higher incidence of complete damage, producing a large variance in anisotropy decays. Consequently, a large variance in higher radiation dose explains more severe destruction of intracellular structures. Anisotropy decays with different extent of radiation damage are simulated in Fig. S11 in the Supplementary Information.

By contrast to the positive correlation between anisotropy decay and radiation dose, no systematic correlation is observed between the flow rates and the anisotropy decay. Further experiments will be necessary to investigate, whether flow rates beyond the maximum one used here (1000 *µ*Lh^−1^, corresponding to 9.26 *µ*m/ms), will diminish the radiation damage on the cell samples. Additionally, it will be interesting to investigate the influence of the scan direction (i.e., in parallel or perpendicular to the flow direction), thus testing if we are able to “outrun” the diffusion of radicals to unexposed regions on the sample or not. The damage to cells from X-ray radiation is a result of both direct and indirect processes (Riley, 1994; Hémonnot & Köster, 2017; Nicolas *et al*., 2019*a*). The indirect processes are typically caused by free radicals generated by water radiolysis (Le Caër, 2011; Stark, 2005). These radicals react with cellular components, and with other radicals (Stark, 2005; Nordberg & Arnér, 2001; Kai *et al*., 2025). In pure water environments, these reactions occur extremely, ranging from picoseconds to nanoseconds (Le Caër, 2011; Loh *et al*., 2020; Riley, 1994). Additionally, radicals are transported through the water very quickly due to the Grotthuss mechanism.

## 4 Summary and conclusion

We demonstrate a method to derive quantitative information from fixed-hydrated biological cells using fast scanning SAXS, despite short exposure times and low SNR. In order to maintain a hydrated state of the cells and ensure a close-to-physiological sample environment, a dedicated microfluidic chamber is employed. This chamber enables stable operation for prolonged periods of time on the order of several hours with a wide range of possible flow rates. In our study, we flush buffer through the chamber, because we study fixed-hydrated samples, however, it is equally well possible to replace the buffer by cell medium when investigating living cells. The collected SAXS data are analyzed using a novel noise filtering method based on optimizing the *q*-range such that the scattering angle with meaningful signal are captured, but as little of noisy signal is included. This approach allows for the reduction of background noise without any observable information loss, as observed from inspecting the dark filed contrast images. The optimized *q*-range also enables us to derive orientation and anisotropy of sub-cellular structures using the circular mean and variance of the azimuthal intensity profile. Comparing this signal with visible light fluorescence images of the keratin bundles and networks in the cells, we observe a strong correlation, confirming that structural information collected using fast scanning SAXS indeed originates from intracellular structures.

Biological cells are very sensitive to radiation damage. Therefore, in order to demonstrate the utility of our approach combining microfluidics and dedicated data analysis for fast scanning SAXS, we perform systematic analyses of radiation-induced damage to intracellular structures. Interestingly, damage to the structures associated with anisotropy occurs already during the first scan, despite the very short exposure time of few ms. Thereafter, the anisotropy signal remains at a low level but does not decay further. Notably, this damage is hardly observed in the dark field contrast images. As expected, the extent of the radiation damage is positively correlated with the radiation dose. However, we do not observe a correlation with the flow rate in the range used in our experiments. Our study demonstrates that state-of-the-art synchrotron radiation with high flux and focused beams allows for the analysis of structural information stemming from intracellular structures, beyond mere cell shape segmentation. Our approach, including the X-ray compatible microfluidic chamber can be extended to living cells in a straightforward manner.

## Acknowledgements

We are grateful for the long term proposal beam time (SC-5436) granted by ESRF (The European Synchrotron, Grenoble, France), and in particular outstanding support by the ID 13 staff and the PSCM (Partnership for Soft Condensed Matter), specifically Pierre Lloria, Peter van der Linden. We also thank Peter Luley for technical support, Peter Gawlitza and Sven Niese for providing MLL-lens prototypes used for these experiments, and Alexander Egner and Tim Salditt for fruitful discussions.

## Funding

This work was financially supported by the German Ministry of Education and Research (BMBF) under grants No. 05K22MG3 and 05K25MGA and the German Research Foundation (DFG) under Germany’s Excellence Strategy - EXC 2067, ‘Multiscale Bioimaging: from Molecular Machines to Networks of Excitable Cells’ (MBExC, grant No. EXC 2067/1-390729940) and in the framework of DAPHNE4NFDI (grant No. 460248799).

### Data availability

Upon acceptance of the manuscript, the data will be uploaded to a repository (GRO.data) and the respective doi will be included here.

