## Supplementary Information for "Fast scanning small angle X-ray scattering of hydrated biological cells"

### Supplementary Material for: Fast scanning small angle X-ray scattering of hydrated biological cells

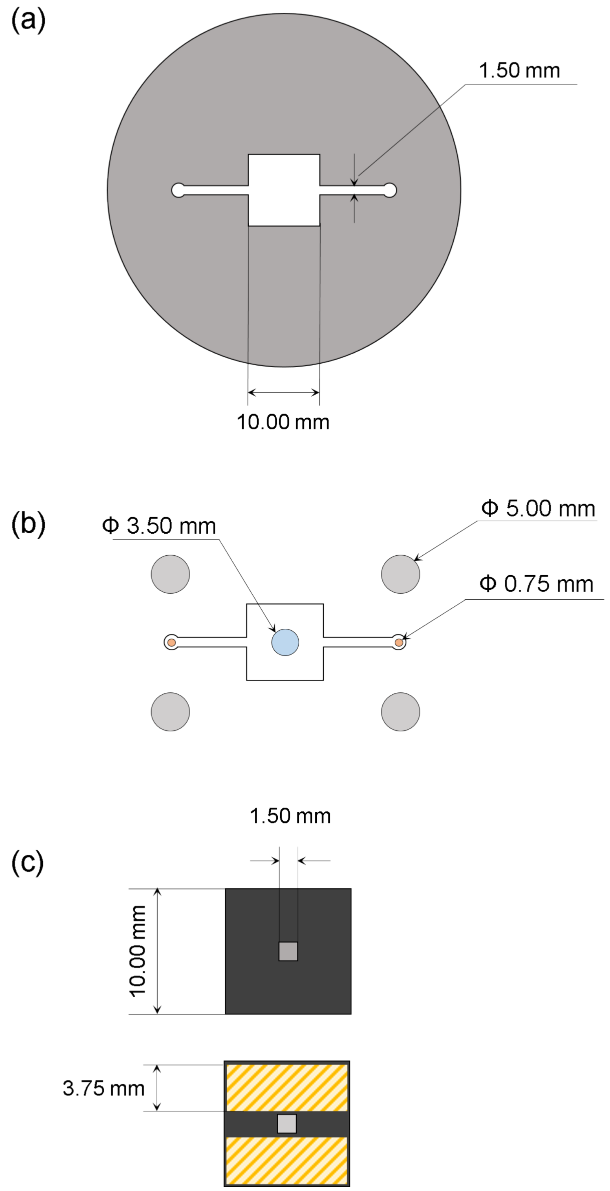


Figure S1: (a) Photomask design. The outer diameter is 2 inches and the structure is positioned in the center. (b) Structure casted in PDMS with punched holes; diameters: 0.75 mm for the inlet and outlet (orange), 3.5 mm for the X-ray observation window (blue), and 5.0 mm for the screws (gray). (c) SiN membranes; the yellow-hatched area on the bottom membrane is the custom-ordered spacer. The thickness of the silicon frame is 200 $\mu$m and the thickness of the SiN window is 1 $\mu$m. The spacer thickness is 20 $\mu$m.

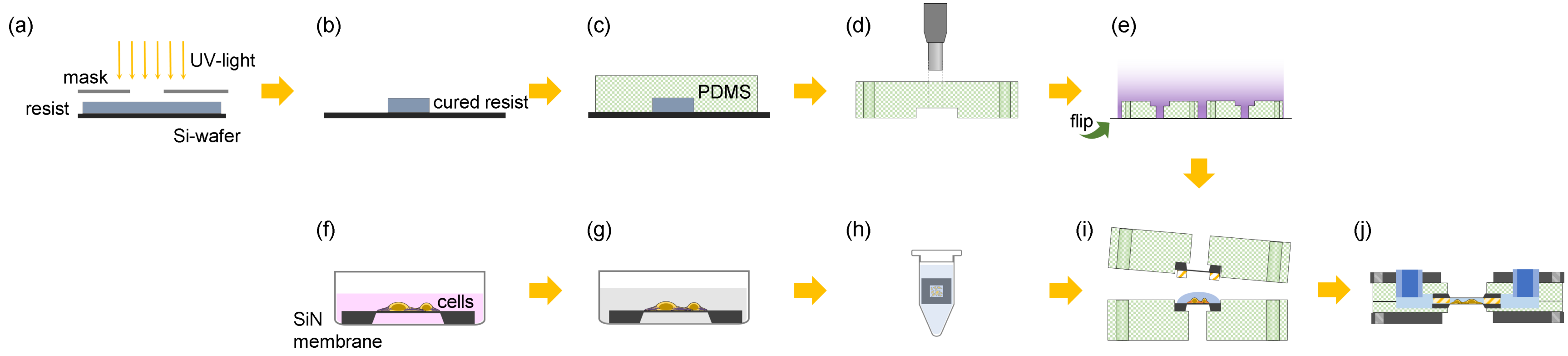


Figure S2: Chamber fabrication process; a to d: photolithography and PDMS replication; f to h: sample preparation process as described in Sec. 2.1 of the main text. (a) A thin layer of resist is spin coated on a Si-wafer and exposed to UV-light through the photomask. (b) After developing, the UV-exposed region in the resist remains and forms the channel structure. (c) PDMS block is cast from the master. (d) Access holes for X-rays, screws, and tubing are punched. (e) Plasma-activation of the surface of the two PDMS blocks. (f) Cells are grown on SiN membranes. (g) Cells are chemically fixed by 4 % formaldehyde and stored in PBS. (h) SiN membrane with cells is stored in PBS. (i) SiN membranes with cells and with SU-8 spacer are placed on each PDMS blocks and sandwiched. (j) Tubings are placed and the metal frames clamp the entire device with screws.

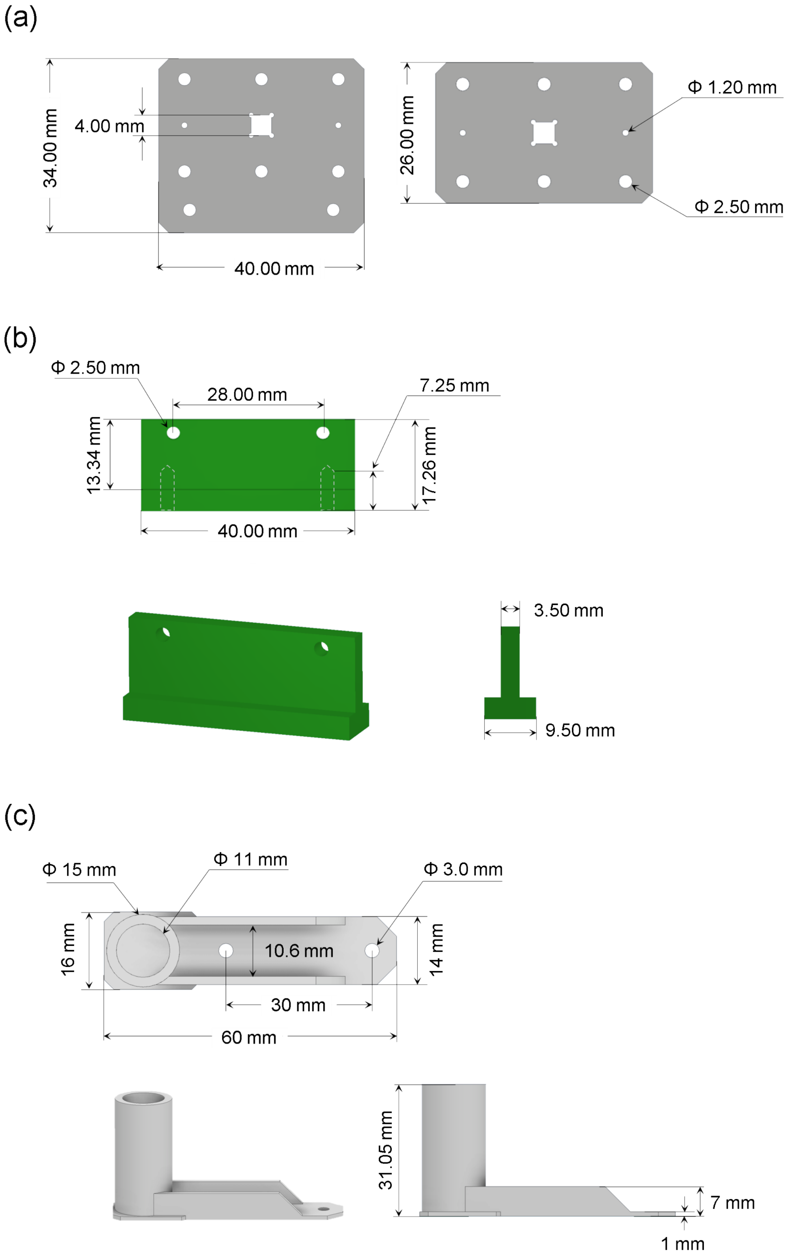


Figure S3: Design of the metal frames and the chamber stand. (a) Metal frames; left: located downstream with an extension to attach it to the chamber stand in b; right: located upstream; holes with a diameter of 2.5 mm are included for clamping using screws; the square holes in the center of both frames constitute the X-ray observation windows. (b) Design of the chamber stand; the two holes on top are used to attach this part to the metal frames shown in a. The thickness of the stand is adequately selected to load the PDMS on it for better stability. The dashed lines illustrate embedded screw holes, which are used for mounting on the sample stage. (c) The design of the waste container is compatible with 1.5 mL or 2 mL reaction tubes. The chamber stand in b is located between the extended wing structures at the bottom. The adapter plate for the piezo sample stage is combined with the stand by two screws, which pass through the waste container.

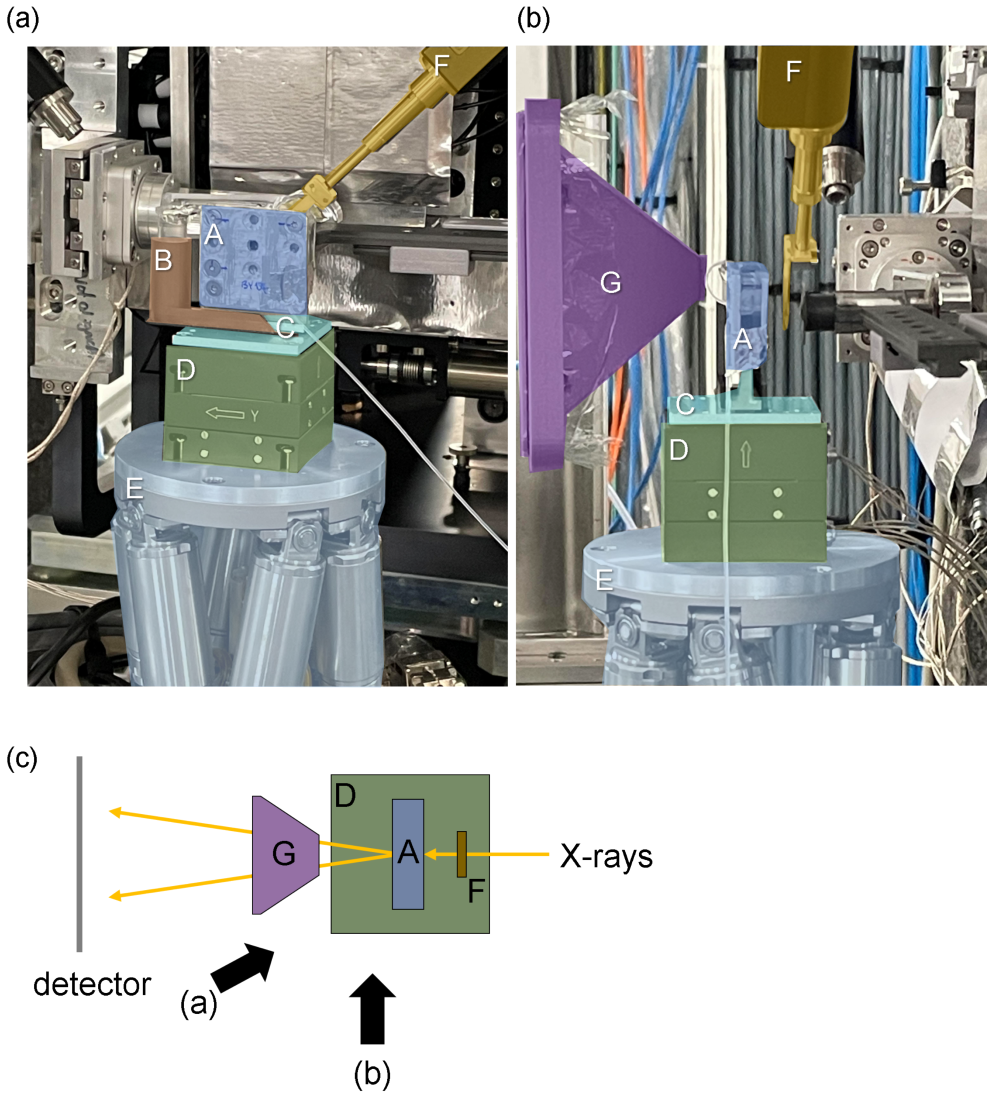


Figure S4: Photographs (a, b) and schematic (c) of the end station with the mounted sample. **A**: sample chamber, visible are the metal frames; **B**: waster container; **C**: chamber stand; **D**: piezo sample stage; **E**: hexapod; **F**: X-ray aperture and holder; **G**: flight tube. (a) and (b) Photographs taken form different angles. (c) Schematic top view; black arrows indicate the observation angles for a and b.

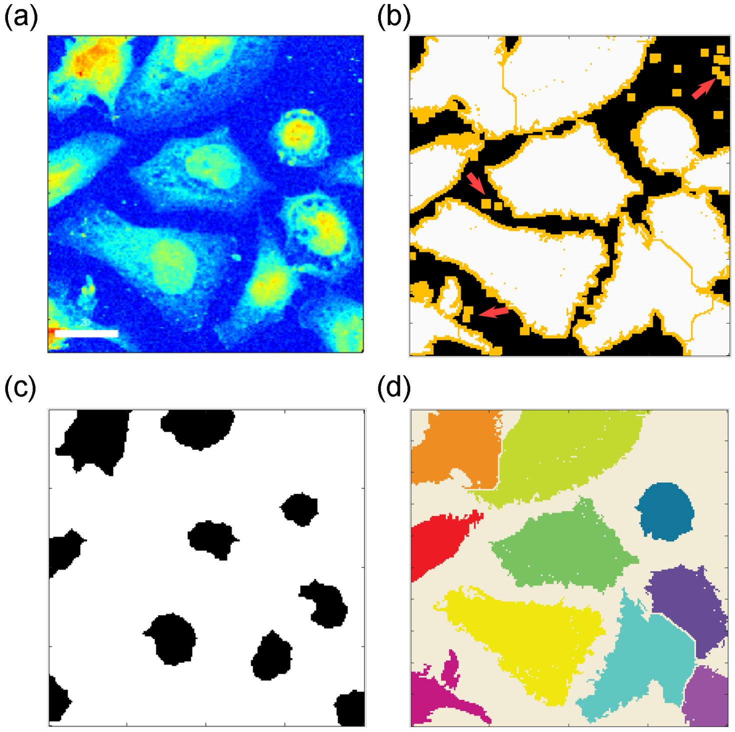


Figure S5: Example of a dark field contrast image and masks generated by different methods. (a) Original dark field contrast image. (b) Background and foreground mask generated by ACSAT. The white regions indicate foreground, and the black regions indicate the background. Added pixels after applying a Gaussian filter are indicated by red arrows. The yellow regions indicate gaps between the background and the foreground after morphological dilation with a circular structure element with a diameter of 3 pixels. (c) Nuclear mask generated by morphological reconstruction. (d) Final cell annotation after applying the marker-controlled watershed method.

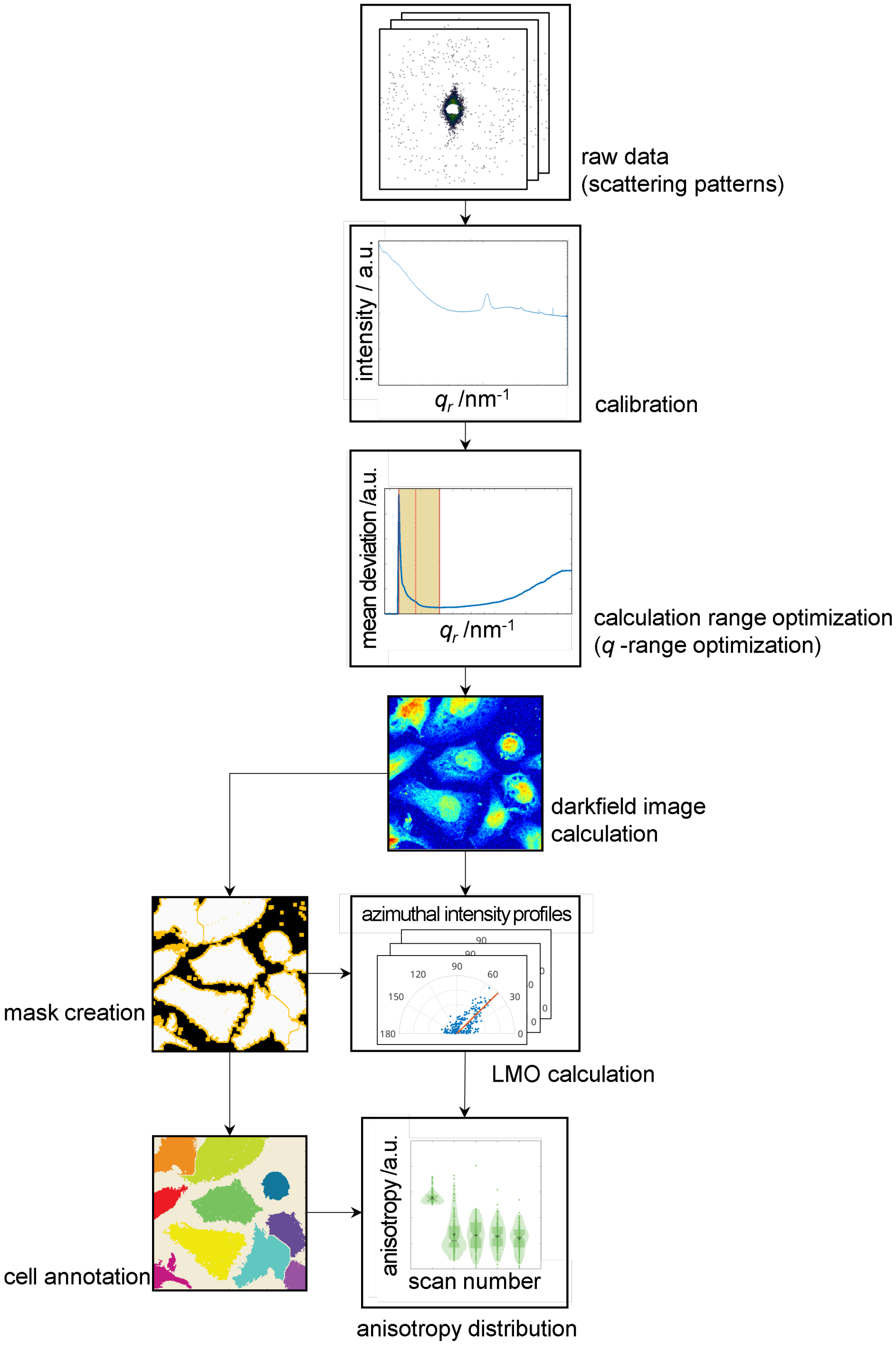


Figure S6: Work flow for the data analysis (further details are described in Sec. 2.4 in the main text): Calibration is performed using AgBeh dataset; the $q$-range is optimized to minimize the noise using a selceted dataset; a dark field image is calculated; the local orientation is determined, including orientation and anisotropy maps; to analyze the anisotropy decays of single cells, cells in a single dark field image are segmented and annotated; the anisotropy distribution of a single cell is represented as one violin plot in the figure.

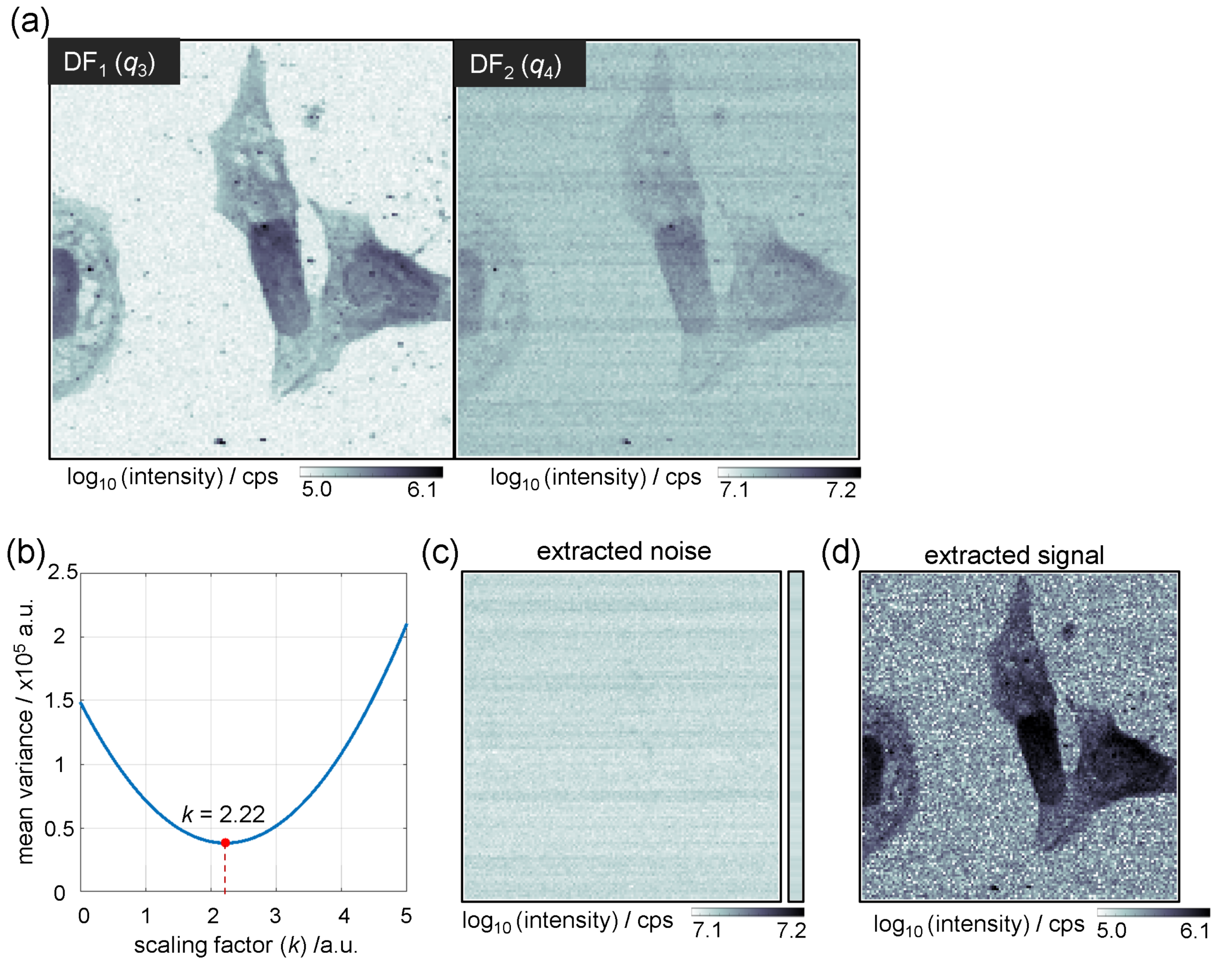


Figure S7: Noise extraction and analysis. (a) Dark field contrast images: The stripe-shaped noise is minimized in $DF_1$, and maximized in $DF_2$, as described in Sec. 3.1 in the main text. Assuming the stripe noise is additive, the cell shapes in $DF_2$ can be considered as a linearly scaled version of $DF_1$ with a scaling parameter of $k$. (b) The mean variance of differences between pixel values in $DF_2$ and $k \times DF_1$ is plotted against $k$; $k$ is determined by the minimum of the curve. (c) $DF_2$ - $k \times DF_1$. Only the stripe pattern is extracted, showing that the noise is indeed additive. The longitudinal box on the right hand side shows the averaged intensity of each row. (d) Noise-subtracted $DF_2$: Averaged intensities are subtracted row by row from $DF_2$. The cell shapes are still visible.

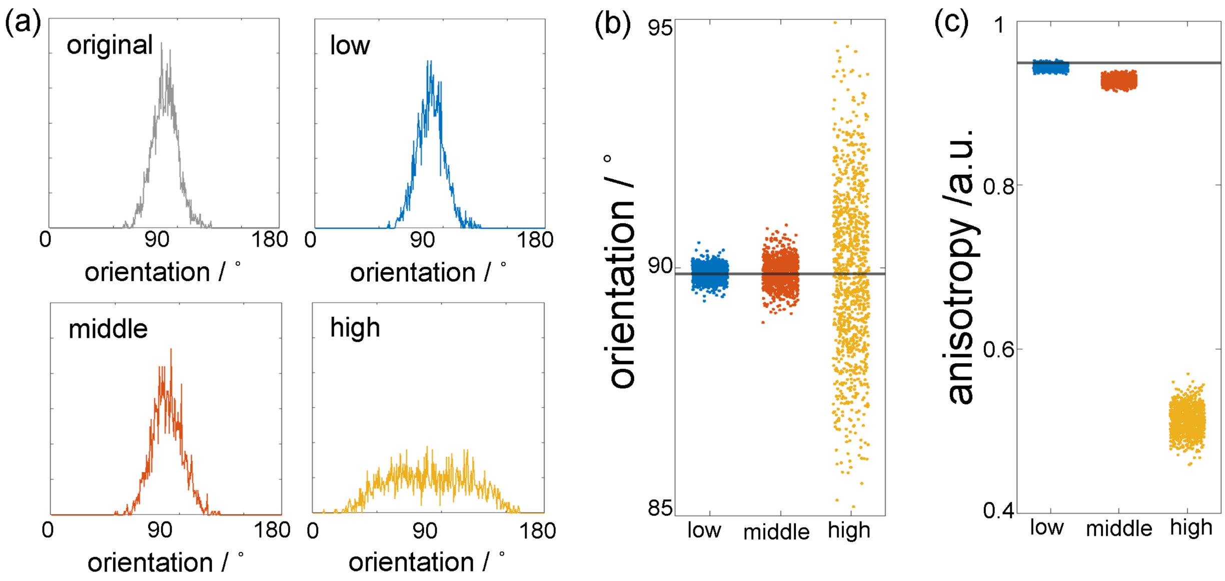


Figure S8: Simulation showing the difference in sensitivity between the orientation and anisotropy. (a) The input signal is a merged data set of two random datasets, each consisting of 1,000 random normally distribution numbers with a mean of 90° or 270°, respectively, and a standard deviation of 10° to obtain a simulated data set similar to the real data. The added noise consists of random integer sets, uniformly distributed in a range of [-5, 5], [-10, 10], and [-50, 50], denoted 'low', 'middle', and 'high', respectively. (b) Corresponding orientation and (c) anisotropy. The black lines show the values of the original signal. The orientation values are distributed around the original dataset's value (90°) with different variances in the noise level, whereas the anisotropy values are shifted towards lower values with larger variance for higher noise.

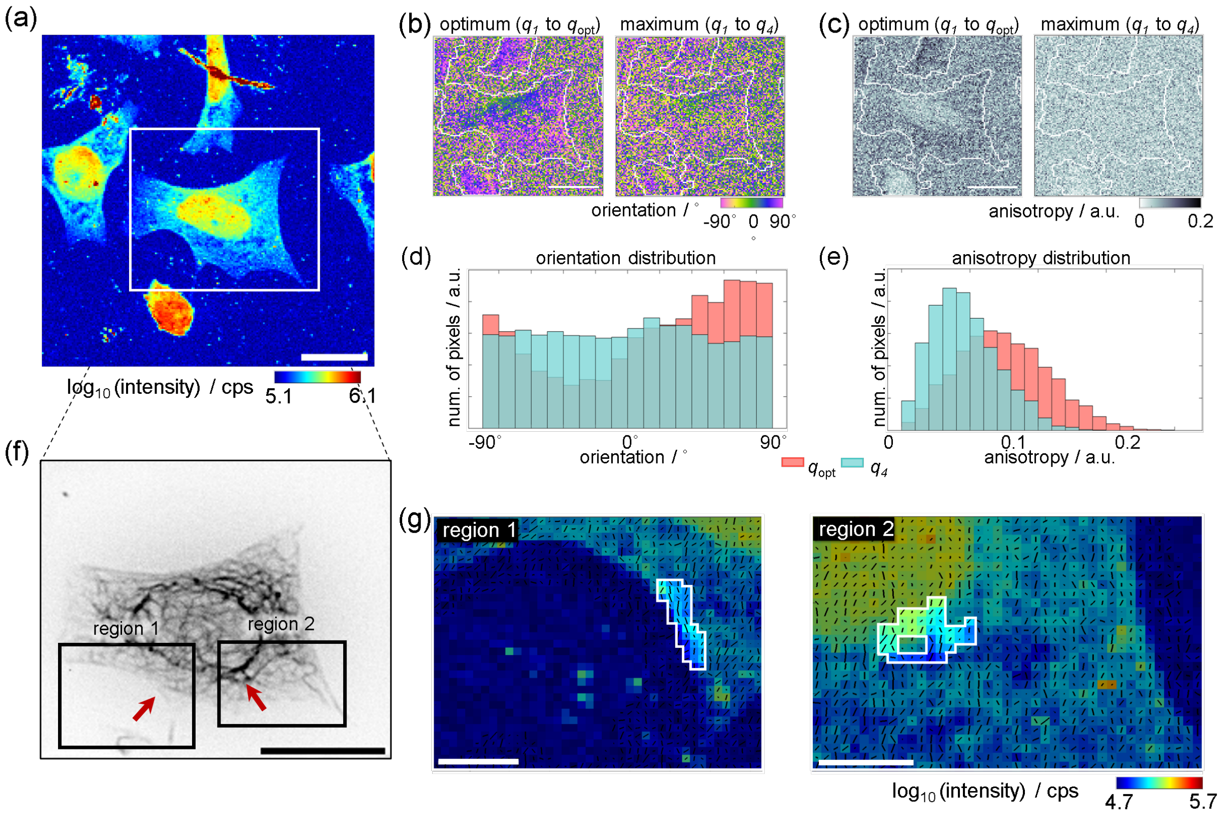


Figure S9: Local orientation analysis (see Fig. 3 in the main text) of an additional data set; the step size is 0.5 × 0.5 $\mu$m and the exposure time is 2 ms. (a) Dark field contrast image calculated using the optimized $q$-range. (b) Orientation maps of the white rectangular region in panel a, calculated using different $q$-ranges: from $q_1$ to $q_{\mathrm{opt}}$ and from $q_1$ to $q_4$. (c) Corresponding anisotropy maps. The white outlines in panel b and c delineate the border of the background area used for the background subtraction. Histograms of (d) orientation and (e) anisotropy distributions of the cell located in the center of panels a-c. (f) Inverted gray scale epifluorescence image of the same cell, acquired prior to the microfluidic chamber assembly. The dark structures are distinct keratin bundles within the cell. (g) Dark field contrast images of the two regions 1 and 2 within the black boxes in panel f, where strongly ordered keratin structures are visible. The black lines in each pixel represent the orientation and anisotropy by their angle and length, respectively. Scale bars in panel a and f: 20 $\mu$m, in panel g: 5 $\mu$m.

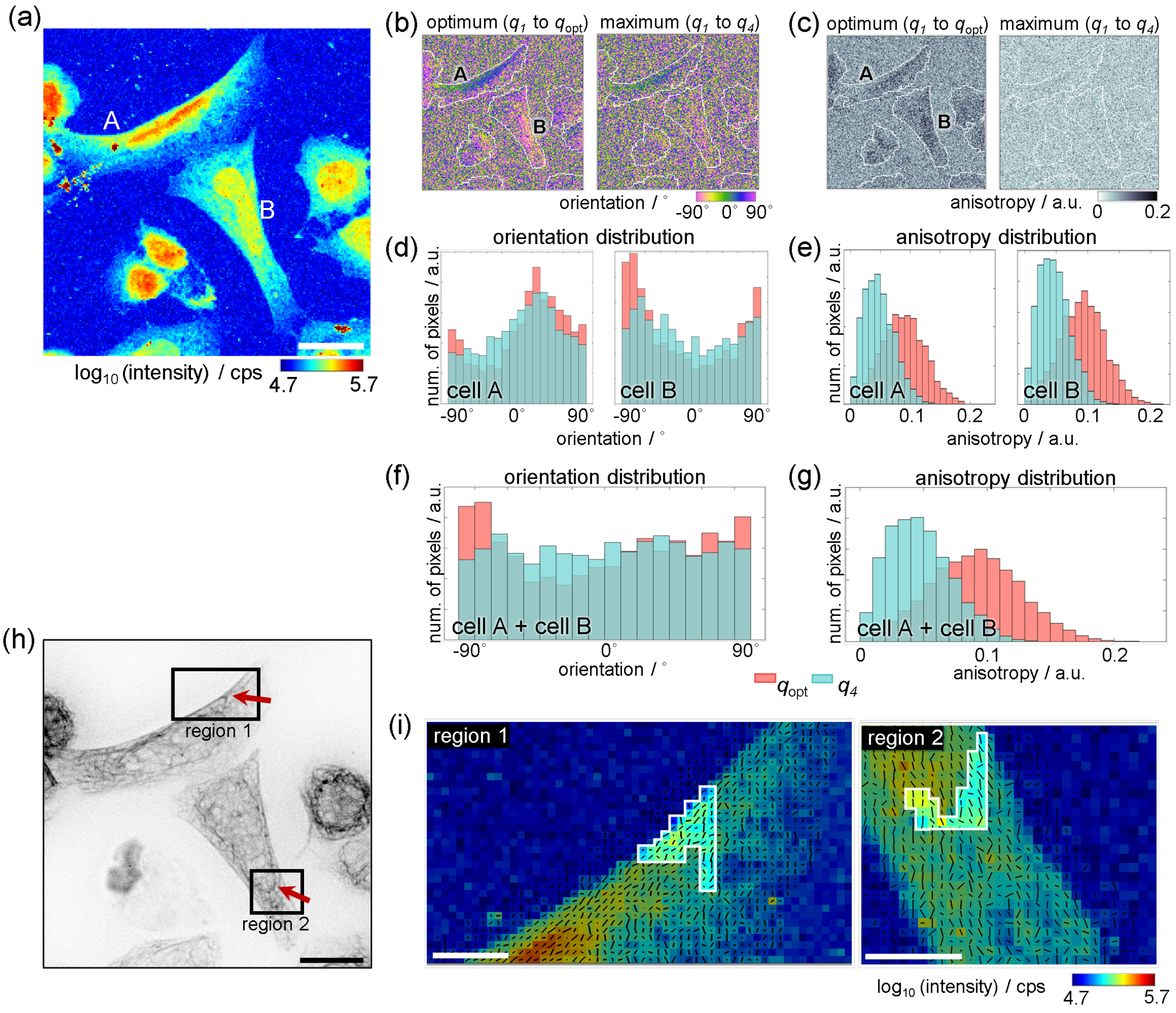


Figure S10: Local orientation analysis (see Fig. 3 in the main text) of an additional data set; the step size is $0.5 \times 0.5$ $\mu$m and the exposure time is 2 ms. (a) Dark field contrast image calculated using the optimized $q$-range. (b) Orientation maps of the region shown in panel a, calculated using different $q$-ranges: from $q_1$ to $q_{\mathrm{opt}}$ and from $q_1$ to $q_4$. (c) Corresponding anisotropy maps. The white outlines in panel b and c delineate the border of the background area used for the background subtraction. Histograms of (d,f) orientation and (e,g) anisotropy distributions of the cells A and B indicated in panels a-c. (h) Inverted gray scale epifluorescence image of the same cells, acquired prior to the microfluidic chamber assembly. The dark structures are distinct keratin bundles within the cells. (i) Dark field contrast images of the two regions 1 and 2 within the black boxes in panel h, where strongly ordered keratin structures are visible. The black lines in each pixel represent the orientation and anisotropy by their angle and length, respectively. Scale bars in panel a and h: 20 $\mu$m, in panel i: 5 $\mu$m.

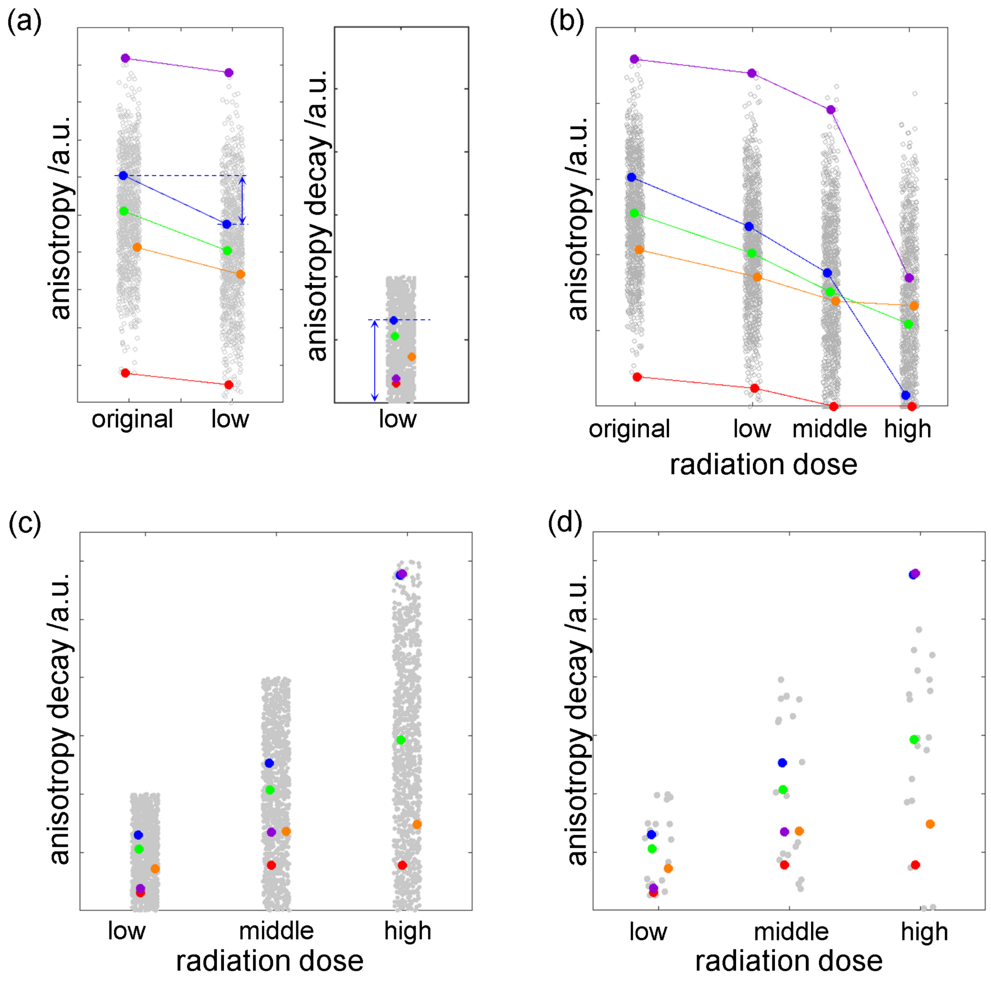


Figure S11: Simulation of anisotropy decay distribution. (a) Explanation of how to determine the anisotropy decay. The original signal consists of 1,000 random normally distributed numbers between [0, 1] with a mean of 0.5 and a standard deviation of 0.1. Each data point indicates an averaged anisotropy value of the cytoplasm region of a single cell; thus the 1,000 data points represent 1,000 cells. The anisotropy decay is the difference between the original and radiation-damaged anisotropy values, where the extent of the radiation damage is randomly selected within a constrained range according to the radiation dose (see blue dashed lines and arrow for an example). Any anisotropy decays smaller than zero are set to zero. In the visualization, five example data points (colored) are randomly selected and tracked. The lengths of the arrows are different in the left and the right plots due to the scaling. (b) Influence of different levels of radiation dose. The extent of radiation damage is set to [0, 0.2] for the low dose, [0, 0.4] for the middle dose, and [0, 0.6] for the high dose. (c) Anisotropy decay values of all 1,000 data points. The higher the dose the larger the variance. (d) 25 random data points, including the previously selected 5 data points, which exhibit a similar trend to Fig. 4 in the main text. This similarity explains that the large variance in the anisotropy decay at high doses indicates more damage.
